# A Major Role for Common Genetic Variation in Anxiety Disorders

**DOI:** 10.1101/203844

**Authors:** Kirstin L. Purves, Jonathan R. I. Coleman, Sandra M. Meier, Christopher Rayner, Katrina A. S. Davis, Rosa Cheesman, Marie Bækvad-Hansen, Anders D. Børglum, Shing Wan Cho, Jürgen Deckert, Héléna A. Gaspar, Jonas Bybjerg-Grauholm, John M. Hettema, Matthew Hotopf, David Hougaard, Christopher Hübel, Carol Kan, Andrew M. McIntosh, Ole Mors, Preben Bo Mortensen, Merete Nordentoft, Thomas Werge, Kristin K. Nicodemus, Manuel Mattheisen, Gerome Breen, Thalia C. Eley

## Abstract

Anxiety disorders are common, complex psychiatric disorders with twin heritabilities of 30-60%. We conducted a genome-wide association study of Lifetime Anxiety Disorder (n = 83 565) and an additional Current Anxiety Symptoms (n= 77 125) analysis. The liability scale common variant heritability estimate for Lifetime Anxiety Disorder was 26%, and for Current Anxiety Symptoms was 31%. Five novel genome-wide significant loci were identified including an intergenic region on chromosome 9 that has previously been associated with neuroticism, and a locus overlapping the *BDNF* receptor gene, *NTRK2*. Anxiety showed significant genetic correlations with depression and insomnia as well as coronary artery disease, mirroring findings from epidemiological studies. We conclude that common genetic variation accounts for a substantive proportion of the genetic architecture underlying anxiety.

## Introduction

Anxiety disorders are amongst the most common classes of psychiatric disorders worldwide^1, 2^ and have a global lifetime prevalence of ∼16%^1^. They were responsible for an estimated 27 121 years lost due to disability globally in 2017, accounting for 3% of the total from all conditions measured^3^. Anxiety disorders aggregate in families, and twin heritability estimates range from 30-60%, dependent upon the participant age and specific trait or disorder being assessed^4, 5^. Although many candidate gene studies of anxiety disorders have been carried out, these associations have not proven robust^6^. As is the case with other psychiatric disorders, such as schizophrenia^7^ and depression^8^, it is likely that a multitude of common genetic variants with modest effects, in addition to environmental factors, underlie the risk for anxiety disorders^6^.

Clinically, anxiety is divided into several sub-classifications, including generalised anxiety disorder, social anxiety disorder, panic disorder, agoraphobia, and specific phobias. However, there is evidence for commonalities across anxiety disorders at both the phenotypic^9^ and genetic^10, 11^ level. The majority of the data indicating genetic overlap between different anxiety traits/disorders comes from the twin literature^10, 12–14^, with clear evidence that the shared genetic component between anxiety disorders is larger than the unique contributions to any one disorder.

Furthermore, the covariance between anxiety disorders and depression is best explained by a single genetic factor with some evidence for additional phobia specific genetic factors^13^. A recent review summarised a range of research strongly indicating that current diagnostic boundaries between anxiety disorders are unlikely to reflect biologically distinct disorders ^6^. Similarly, several twin studies have found high genetic correlations between the personality trait of neuroticism and anxiety disorders (including generalised anxiety disorder^15–17^, social and situational phobias and agoraphobia^17, 18^). As such, it is likely there is a single underlying liability distribution, with variation in anxiety related traits occurring to different degrees across the population, driven in part by common genetic variants^10, 13, 19, 20^. If this were the case, it is likely that anxiety disorders share a common polygenic influence, which may explain the shared phenotypic and genetic structure identified in the twin literature.

The detection of genetic variants and genes associated with anxiety disorders has made slow progress, limited by small sample sizes. There have been some suggestive hits including *TMEM132D* for panic disorders, but few have been replicated^6^. A meta-analysis of genome-wide association studies of several anxiety disorders (N= 18 186) identified one genome-wide significant locus with anxiety caseness (n_cases_ = 3 695; n_controls_ = 13 615) and a second with a quantitative factor score of broad anxiety^11^. Finally, a SNP within *RBFOX1* was found to be significantly associated with anxiety sensitivity in a small cohort of twins^21^. The proportion of variation in generalised anxiety disorder symptoms explained by individual genetic variation (SNP heritability; h^2^_SNP_) was estimated at 7.2% in a modest sample (n=12 282) of Hispanic/Latino adults^22^, and 14% in the meta-analysis described above, which consisted largely of individuals with European ancestry^11^.

To date, none of these findings have been replicated, although *RBFOX1* was significantly associated with major depression in a recent genome-wide association analysis^8^. Analyses of significantly larger samples are required to further our understanding of the underlying genetic architecture of anxiety disorders and current anxiety symptoms at the population level. With this aim, we conducted genome-wide association studies (GWAS) of two anxiety phenotypes: Lifetime Anxiety Disorder combining self-reported clinical lifetime diagnosis of an anxiety disorder and probable DSM-IV generalised anxiety disorder; and Current Anxiety Symptoms, counting as cases anyone who reported at least moderate symptoms of generalised anxiety disorder in the two weeks preceding assessment. Data were drawn from the UK Biobank, a large population sample. Two cohorts containing anxiety disorder cases and controls were considered for the purposes of replication; the Anxiety NeuroGenetics Study (ANGST)^11^, and a sample from the Danish iPSYCH study^23^. Further replication-extension analyses were undertaken using summary statistics from two highly relevant phenotypes – Neuroticism (in a distinct subset of the UK Biobank), and major depressive disorder (in a subset of the Psychiatric Genetics Consortium Major Depressive Disorder sample)^8^.

## Methods

### Sample and phenotype definition

Samples were a subset of 126 443 (age 46-80) individuals of Western European ancestry who took part in the UK Biobank online mental health follow-up questionnaire.

#### (1) Self-report or probable lifetime anxiety disorder (Lifetime Anxiety Disorder)

This was our primary phenotype. Cases met one of two definitions. First was self-reporting a lifetime professional diagnosis of one of the core five anxiety disorders, (generalised anxiety disorder, social phobia, panic disorder, agoraphobia or specific phobia; n=21 108). Further case were defined as meeting criteria for a likely lifetime diagnosis of DSM-IV generalised anxiety disorder based on anxiety questions from the Composite International Diagnostic Interview (CIDI) Short-form questionnaire^24^ (n=4 345). We excluded individuals self-reporting a lifetime diagnosis of schizophrenia, bipolar disorder, autistic spectrum disorder, attention deficit hyperactivity disorder, or eating disorder. Individuals meeting one or other of these criteria resulted in a total of 25,453 cases.

#### (2) Moderate/severe current generalised anxiety symptoms (Current Anxiety Symptoms)

For this secondary phenotype, participants were defined as cases if they obtained a total score on a screening measure for anxiety symptoms over the past two weeks (GAD-7) of >= 10 out of a total score of 21. This is a standard threshold for recent moderate to severe generalised anxiety symptoms as measured by the GAD-7^25^ (n=19 012).

#### Healthy controls

A common control group was identified consisting of a set of screened healthy individuals who did not meet criteria for any mental health disorder or known substance abuse and were not prescribed medication for any psychiatric disorder (n=58 113).

See **Supplementary Information** for more detailed description of each category.

### Independent anxiety-related samples

Data from four other studies were used to replicate and extend our analyses. These included GWAS summary statistics from the Anxiety NeuroGenetics Study^11^ of individuals meeting DSM criteria for one of the five core anxiety disorders (generalised anxiety disorder, social phobia, panic disorder, agoraphobia or specific phobia (n=3 695); and controls (n=13 615). In addition to the case-control phenotype, for some analyses we also report the ANGST factor score derived from combining controls and individuals reporting full and sub-threshold symptoms of the five anxiety disorders; (n=18 186) ^11^. Second, summary statistics from an unpublished subset of lifetime anxiety cases (n=2 829) and controls (n=10 386) from the iPSYCH cohort in Denmark^23, 26^, defined using the same inclusion and exclusion criteria as for our primary phenotype. Third were summary statistics from an analysis of total neuroticism score in all individuals who completed a baseline assessment in the UK Biobank (UKB Neuroticism), who were not included as either a case or control in the Lifetime Anxiety Disorder phenotype (n=241 883). Finally we used summary statistics for Major Depressive Disorder from the most recent Psychiatric Genomics Consortium depression analysis (PGC MDD)^8^, excluding samples drawn from the UK Biobank or 23&Me (n_cases_=45 591; n_controls_=97 674).

In summary, anxiety-related phenotypes from four replication samples were available for comparison with the core UK Biobank Lifetime Anxiety Disorder phenotype; 1) ANGST Anxiety Disorder, 2) iPSYCH Anxiety Disorder, 3) UKB Neuroticism, and 4) PGC MDD. See **Supplementary Table 1 f**or sample sizes for each phenotype.

### Genotyping and quality control

Genotype data were collected and processed as part of the UK Biobank extraction and quality control pipeline^27^. Only SNPs imputed to the Haplotype Reference Consortium (HRC) reference panel or genotyped SNPs were used for these analyses. SNPs with a minor allele frequency >0.01 and INFO score >0.4 (indicating well imputed variants) were retained. For details see **Supplementary Information.**

## Statistical Analyses

### Primary analyses

#### Genome-wide association analyses

Analyses were limited to individuals of European ancestry defined by 4-means clustering on the first two ancestry principal components. Covariates (age, sex, genotyping batch, assessment centre, and the first six genetic principal components) were regressed out of each phenotype using logistic regression^28^. Resulting residuals were used as dependent variables in three linear genome-wide association analyses, using BGENIE v1.2 software^27^. Variants surpassing genome-wide significance (p<5×10^−8^) were annotated using Region Annotator software to annotate known protein coding genes within the regions of significance (https://github.com/ivankosmos/RegionAnnotator).

#### SNP heritability

To estimate the proportion of variance explained by common genetic variants (h^2^_SNP_) for the two UK Biobank anxiety phenotypes we used variance component analyses conducted in BOLT-LMM^29^. Estimates (and standard errors) were converted to the liability scale assuming: (1) accurate sampling rate in the UK Biobank (sample prevalence = population prevalence), (2) sample prevalence < population prevalence; and (3) sample prevalence > population prevalence (**Supplementary Table 2**).

#### Genetic correlations

LD score regression^30^ was used to estimate genetic correlations between 1) UK Biobank LIfetime Anxiety Disorder, Current Anxiety Symptoms and all replication samples, and 2) UK Biobank Lifetime Anxiety Disorder and other previously published phenotypes. Genetic correlations are considered significant at p<0.0002, Bonferroni corrected for 251 independent tests.

### Secondary analyses

Further analyses were carried out on the core Lifetime Anxiety Disorder phenotype except where otherwise specified.

#### Partitioned heritability

Stratified LD score regression was used to partition SNP heritability by functional genomic categories. See **Supplementary Information** for results of this analysis.

#### Gene-wise analysis

Gene-wise analyses were computed on GWA summary statistics by MAGMA^31^. See **Supplementary Information** for additional details.

#### Polygenic prediction of anxiety

We examined the extent to which genome-wide polygenic signal from independent SNPs identified in the Lifetime Anxiety Disorder GWAS account for variance in liability to Lifetime Anxiety Disorder case status using a leave-one-out approach. All participants in the Lifetime Anxiety Disorder analysis were randomly divided into ten subgroups of approximately equal size with case: control ratio comparable to the primary analysis. Ten new genome-wide association analyses, with the same controls as in the main analyses, were performed using PLINK 1.9^32^. Each sub group was left out of one analysis and served as the target sample for polygenic scoring. An average R^2^ was derived across all ten iterations of the polygenic scoring at the p-threshold that was best predictive. For detail of polygenic score creation see **Supplementary information.**

#### Testing the dimensionality of anxiety

To consider the degree to which differing levels of severity in anxiety traits reflect the same underlying genetic factors, three independent groups were identified using their current anxiety (GAD-7) symptom scores, including mild, (scoring 5-10, n_cases_=49 144), moderate (10-15, n_cases_=13 788) and severe (>15, n_cases_=5 093). See **Supplementary Table 3** for sample sizes for each analysis. Genetic correlation between these three groups was calculated using LD Score regression^30^.

#### Replication of Significant SNPs

Any SNPs that surpassed genome-wide significance (5×10^−8^) in the core UK Biobank Lifetime Anxiety Disorder analysis were examined for direction and significance of effect in the four replication/extension samples.

#### Meta-analysis

The three Lifetime Anxiety Disorder samples; UK Biobank Lifetime Anxiety Disorder, ANGST Anxiety Disorder and iPSYCH Anxiety Disorder were combined using inverse-variance weighted, fixed-effect meta-analyses in METAL^33^. Only SNPs shared across all three samples were included in this meta-analysis.

### Code availability

Code for all analyses are available at https://github.com/klpurves/UKBB_Anxiety.

## Results

### Genome-wide association

Results of the genome-wide association analyses for the two anxiety phenotypes (Lifetime Anxiety Disorder and Current Anxiety Symptoms) in UK Biobank are shown in Figure 1. Manhattan and Q-Q plots are shown for each phenotype. **Supplementary Table 4** shows the results of region annotation for regions that surpassed genome-wide significance (P <5×10^−8^).

**Figure 1.**
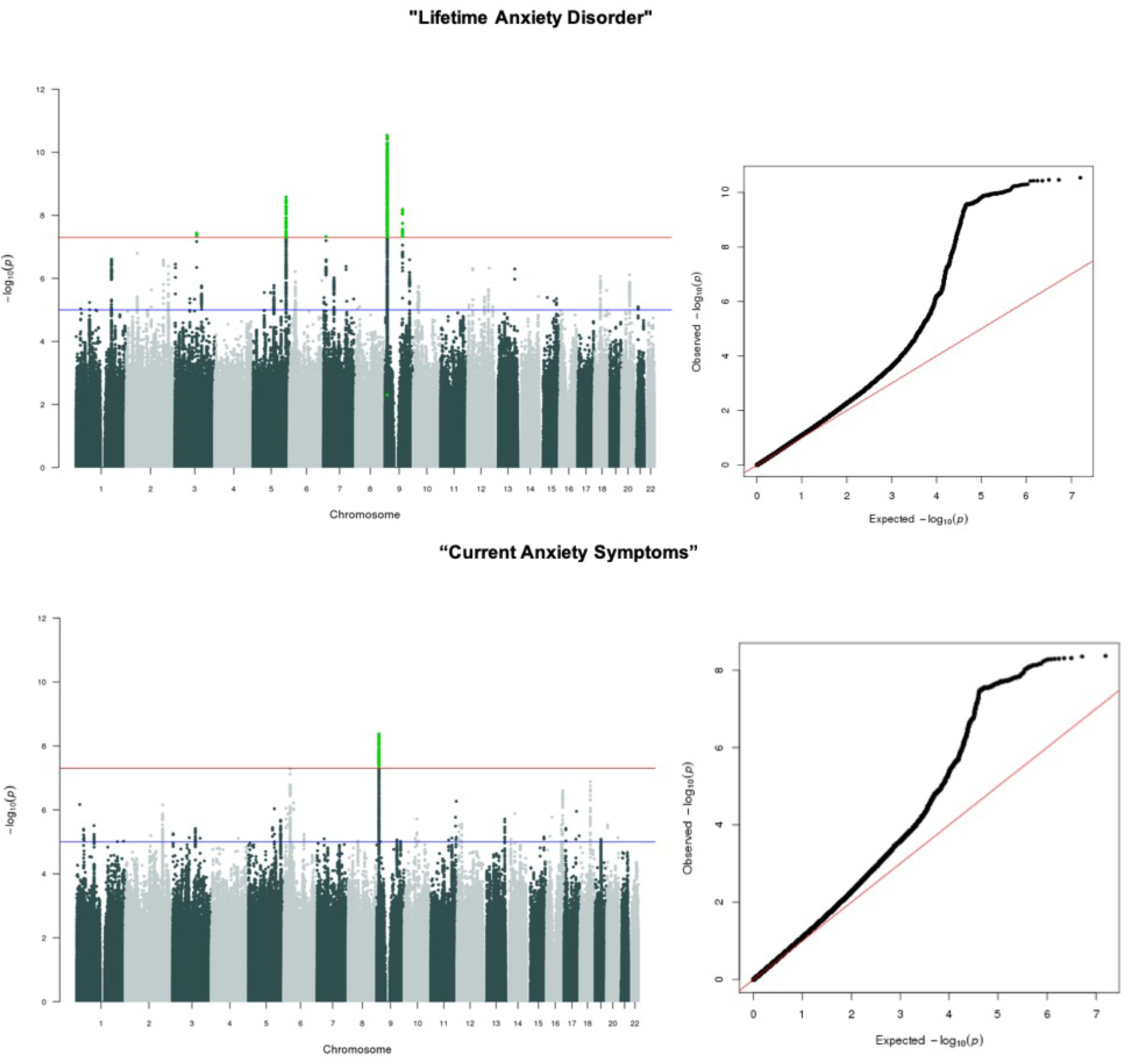
Manhattan and Q-Q plots for UK Biobank anxiety analyses. Figure 1 shows Manhattan and Q-Q plots of p-values of Single Nucleotide Polymorphisms (SNP) based association analyses of anxiety in the UK Biobank. In the Manhattan plots, the threshold for genome-wide significance (P < 5×10^−08^) Is indicated by the red line, while the blue line indicates the threshold for suggestive significance (P < 1×10^−5^)

#### Lifetime Anxiety Disorder

Five regions were significant at the genome-wide threshold of 5×10^−8^ on chromosomes 9 (9p23 and 9q21.33), 7 (7q21.1), 5 (5q15) and 3 (3q11.2). The index SNP for the most significant region on 9p23 was rs10809485 (p=1.6×10^−12^) which is in an intergenic region. The index SNP within 9q21.33 is rs1187280 (p=5.2×10^−8^), which is in an intron for the protein coding gene *Neurotrophic Receptor Tyrosine Kinase 2* (*NTRK2*). The index SNP of the chromosome 7 locus (rs3807866, p=4.8×10^−8^) is within *Transmembrane Protein 106B (TMEM106B)*. The chromosome 5 locus (rs2861139, p=2.6×10^−9^) is in an intergenic region, and the chromosome 3 locus (rs4855559, p=3.7×10^−8^) is in the intron for *Myosin Heavy Chain 15 (MYH15)*. See **Supplementary Figures 1-5** for region plots.

#### Current Anxiety Symptoms

A single genome-wide significant locus was associated with our secondary phenotype in the intergenic region on chromosome 9p23, also associated with Lifetime Anxiety Disorder (r^2^=1)^34^. See **Supplementary Figure 6** for the region plot.

### SNP heritability

Estimates of SNP heritability (h^2^_SNP_) converted to the liability threshold are 0.26 *(SE=0.011)*, and 0.31 *(SE=0.011), for* Lifetime Anxiety Disorder and Current Anxiety Symptoms assuming sample prevalence of 0.20 and 0.18 respectively. **Supplementary Table 5** shows the estimates using different assumptions about population prevalence rates.

### Genetic correlation between independent anxiety-related phenotypes

UK Biobank anxiety phenotypes have high genetic correlations with each other (r_G_ 0.93), and with the ANGST case control phenotype (r_G_ 0.74±0.01), UK Biobank neuroticism (r_G_ 0.73±0.04) and PGC MDD (r_G_ 0.78±-0.10). UK biobank anxiety correlates moderately with the ANGST factor score (r_G_ 0.49±-0.03). Note that it was not possible to compare genetic correlations with the iPSYCH sample, as the estimate of h^2^_SNP_ in that sample did not significantly differ from zero. See **Supplementary Table 6** for genetic correlation and standard error between the UK Biobank lifetime and all other phenotypes.

### Genetic correlations with other traits

Significant genetic correlations between Lifetime Anxiety Disorder and a range of other previously published phenotypes traits are presented in Figure 2. Genetic correlations with Current Anxiety Symptoms are in **Supplementary Table 7.**

**Figure 2.**
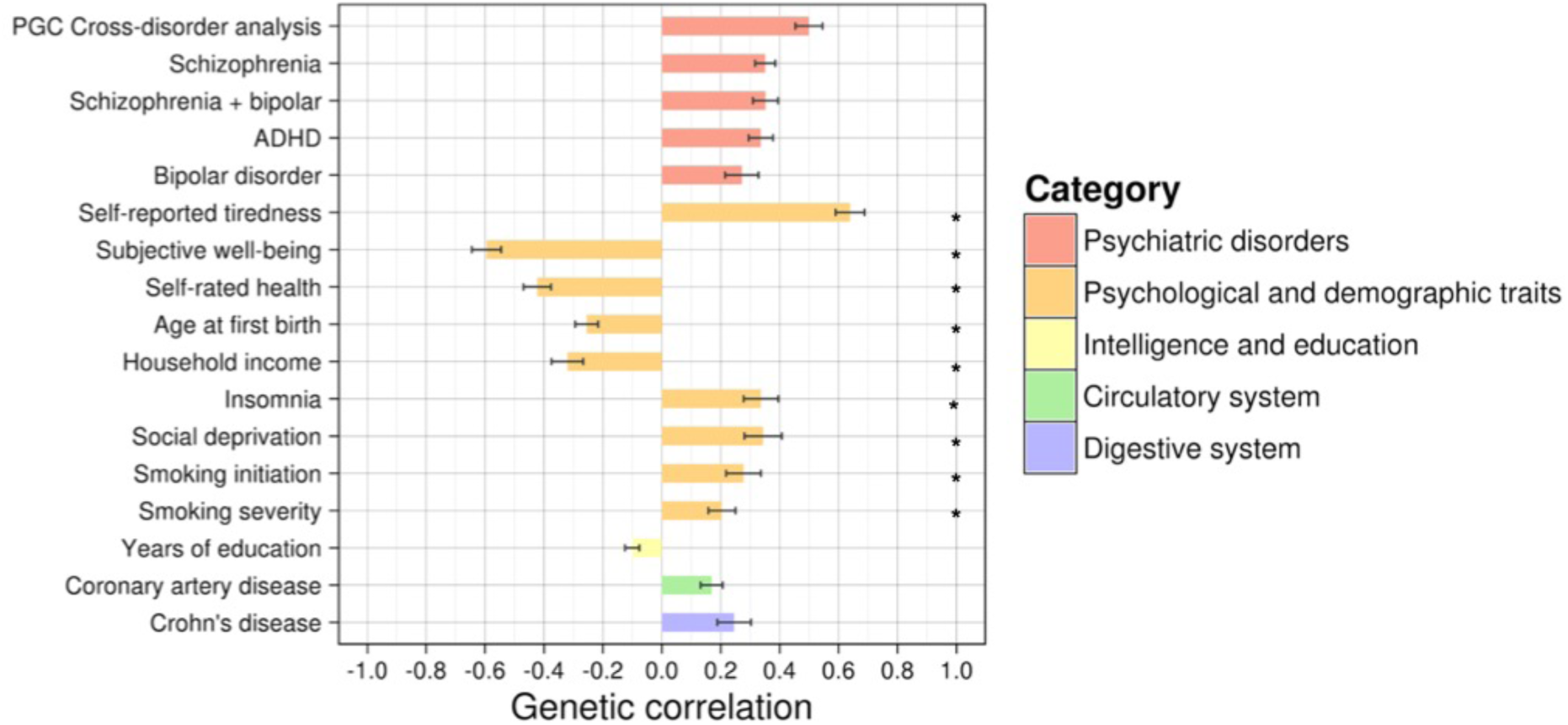
Genetic correlations between UK Biobank Lifetime Anxiety Disorder and significantly associated external traits. Figure 2 shows genetic correlations (r_G_) between UK Biobank “Lifetime Anxiety Disorder” and external phenotypes generated from LDhub. Only traits where correlation exceeded Bonferroni corrected threshold of p < 0.0002 shown here. Depression and Neuroticism were not included in this analysis as they have been included as replication samples in subsequent analyses. * Indicates source study sample wholly or partially included UK Biobank participants. Correlations ordered first by rG within category. Bars show standard error of the estimate.

### Gene-wise analysis

For gene wise associations, significance was determined to be p<2.80×10^−6^ (i.e. 0.05/17831) to account for the number of comparisons. **Supplementary Table 8** shows all gene-wise associations that are significantly associated with Lifetime Anxiety Disorder. Figure 3 presents a Manhattan plot showing the top gene-wise associations with Lifetime Anxiety Disorder per chromosome. The most significantly associated gene in this analysis was the *Transmembrane Protein 106B* (*TMEM106B*; p=6.28×10^−10^).

**Figure 3.**
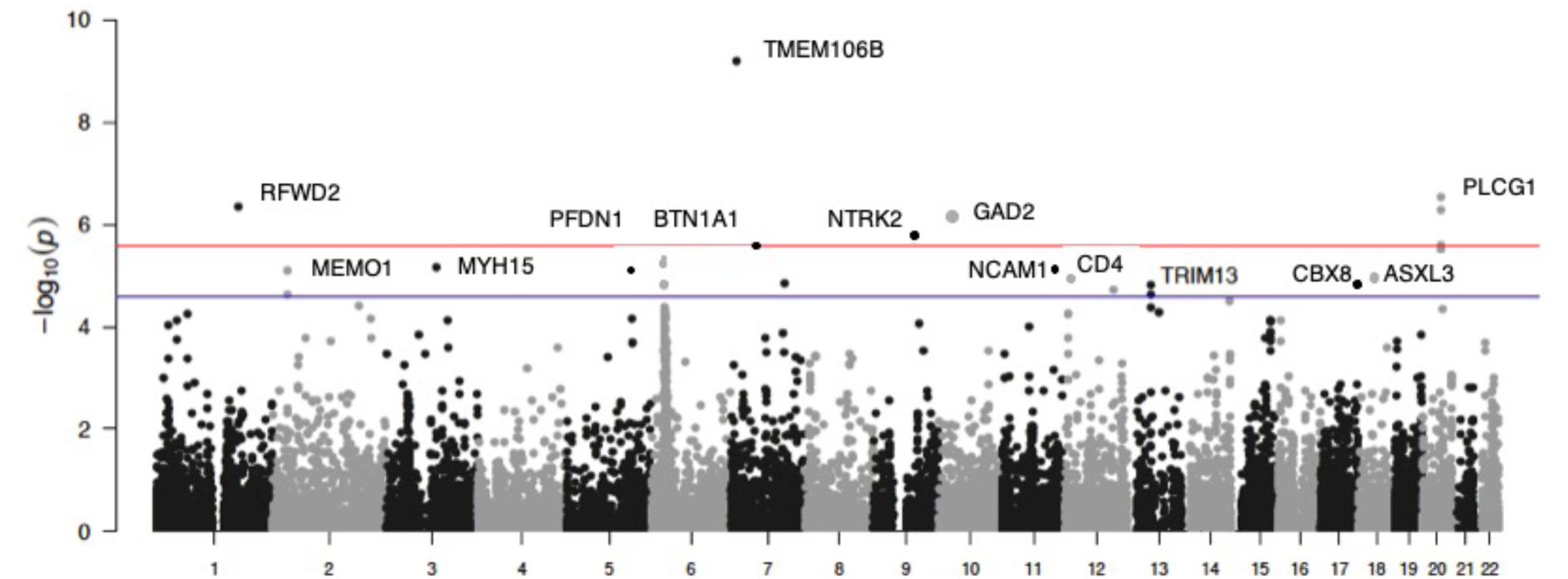
Gene Manhattan plot with top significant gene per chromosome. Figure 3 shows Manhattan plots of p-values of gene-wise association analyses of “Lifetime Anxiety Disorder” in the UK Biobank. In the Manhattan plots, the threshold for gene-wide significance (P < 3×10-06) Is indicated by the red line, while the blue line indicates the threshold for suggestive significance (P < 1×10-5)

### Polygenic prediction of anxiety

Across all ten iterations, a higher polygenic score was significantly associated with increased likelihood of self-reporting Lifetime Anxiety Disorder, explaining on average 0.5% of the variance in outcome on the liability scale across all samples. The relatively low variance explained by the polygenic association with Lifetime Anxiety Disorder relative to our high estimates of h^2^_SNP_ is consistent with what would be expected for a polygenic trait^35^. See **Supplementary Tables 9 – 12** for detailed results.

### Testing the dimensionality of anxiety

Severe current GAD symptoms showed genetic correlations of .76 (*SE*=.08) and .98 (*SE*=.08) with mild and moderate symptoms respectively. Genetic correlation between mild and moderate symptoms was 0.82 (se=.05). See **Supplementary Table 13 and 14** for SNP heritability estimates of these phenotypes and genetic correlations with other anxiety phenotypes respectively.

### Replication and meta-analyses

#### Replication of Significant SNPs

Table 1 shows the p-value and direction of effect for all SNPs found to be genome-wide significant in our core Lifetime Anxiety Disorder analysis, and in the analysis from all four external samples. The locus at 9q21.33 (Index SNP rs10959883) is formally replicated (i.e. significant correcting for 20 independent tests, p<5.25×10^−3^) in both the Neuroticism and MDD samples and has an effect in the same direction in all four samples. The locus annotated to the *NTRK2* (index SNP rs1187280) gene has the same direction of effect in iPSYCH Lifetime Anxiety Disorder and in both Neuroticism and PGC MDD. The locus annotated to the *TMEM106B* (index SNP rs3807866) formally replicates in Neuroticism, and the same direction of effect in iPSYCH Lifetime Anxiety Disorder, and Depression. The locus annotated to the *MYH15* gene (index SNP rs4855559) shows the same direction of effect in the two Lifetime Anxiety Disorder samples, but not in the Neuroticism or Depression analyses.

**Table 1.**
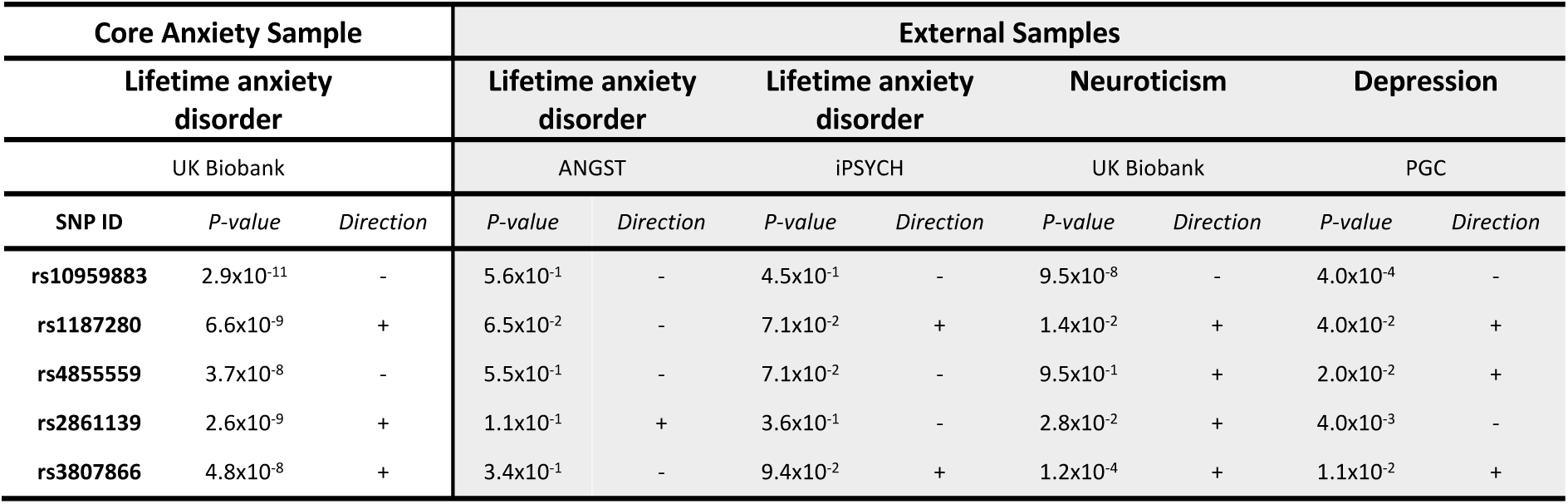
Comparison of genome-wide significant SNPs in “Lifetime Anxiety Disorder” analysis in external samples. Table 1 shows the lead SNP for loci found to be genome-wide significant (P<5×10-8) in UK Biobank “Lifetime Anxiety Disorder” analysis, with p-value and direction of effect in all external samples. External samples include two additional Lifetime Anxiety Disorder analyses; Anxiety NeuroGenetics Study (ANGST)^11^ and Initiative for Integrative Psychiatric Research (iPSYCH) cohorts, and two analyses of related phenotypes; Neuroticism in the UK Biobank (excluding any individuals included in our core “Lifetime Anxiety Disorder” sample) and major depressive disorder from the Psychiatric Genomic Consortium (PGC; excluding samples drawn from UK Biobank and 23&Me)^8^.

#### Meta-analysis

Next, we meta-analysed the core UK biobank Lifetime Anxiety Disorder analysis with the two external Lifetime Anxiety Disorder samples (ANGST and iPSYCH), a combined sample of 114 019 (31 977 cases and 82 114 controls). See Figure 4 for Manhattan and Q-Q plots. Two loci were genome-wide significant. The region on chromosome 9q23 (rs10959577) is nearly identical to the region associated with Lifetime Anxiety Disorder and Current Anxiety Symptoms in the UK Biobank alone (the lead SNPs are ∼70 kb apart with an r^2^=0.27 between them). The second region on chromosome 5 (rs7723509) falls in an intergenic region. See **Supplementary Figure 7** for Manhattan and Q-Q plots for leave-one-out meta-analyses of the UK Biobank, ANGST and iPSYCH samples.

**Figure 4.**
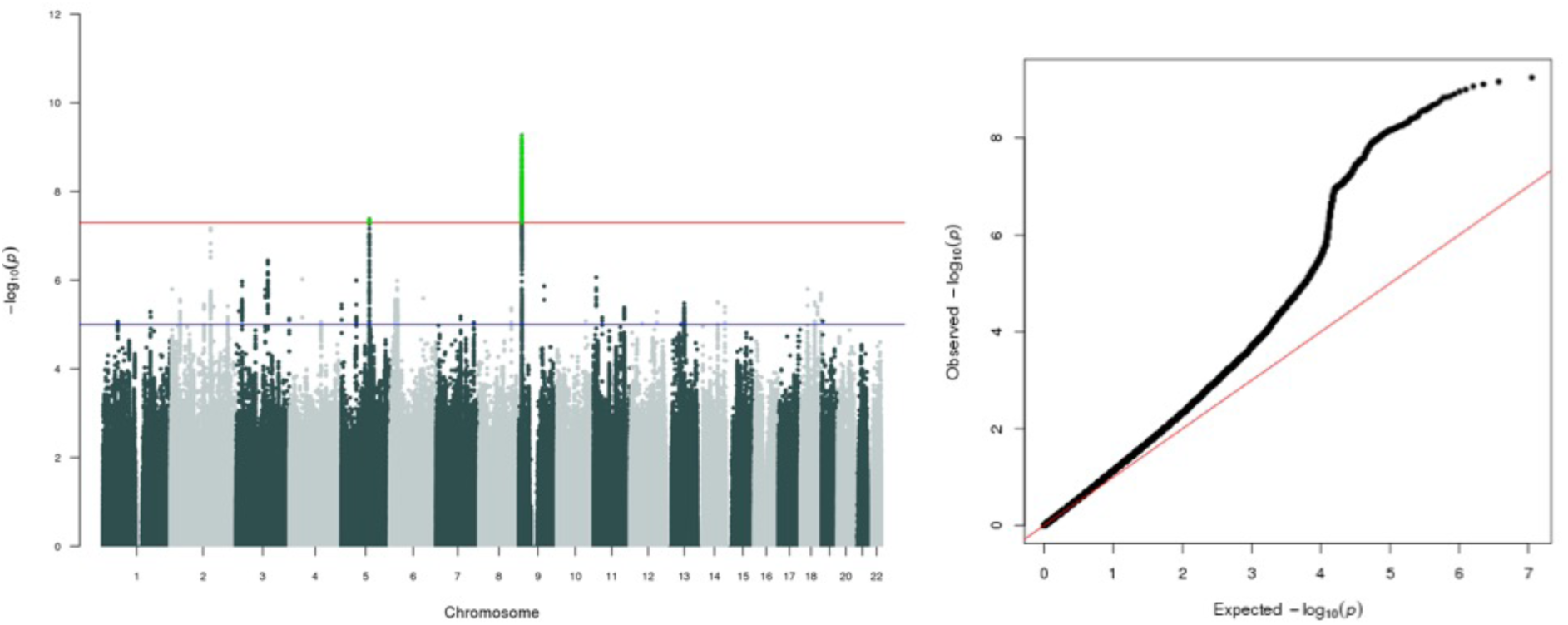
Manhattan and Q-Q plots for meta-analysis of all Lifetime Anxiety Disorder samples. Figure 4 shows Manhattan and Q-Q plots of p-values of Single Nucleotide Polymorphisms (SNP) based association analyses of combined sample meta-analysis across three Lifetime Anxiety Disorder cohorts (UK Biobank, ANGST and iPSYCH). In the Manhattan plots, the threshold for genome-wide significance (P < 5×10-08) Is indicated by the red line, while the blue line indicates the threshold for suggestive significance (P < 1×10-5)

## Discussion

To our knowledge this is the largest GWAS on anxiety conducted, and our core analysis identified five genome-wide significant loci. The first locus is an intergenic region on chromosome 9 previously associated with neuroticism^36, 37^ and depression^8^. The second novel locus on chromosome 9 was in *NTRK2*, a *Brain Derived Neurotrophic Factor (BDNF)* receptor. Together *NTRK2* and *BDNF* regulate both short-term synaptic functions and long-term potentiation of brain synapses (OMIM *600456; https://www.omim.org/entry/600456). Given this key role in brain function, *NTRK2* has been widely investigated in a range of neuropsychiatric traits and disorders^38–46^. The third locus, on chromosome 7, is in *Transmembrane Protein 106B*, a gene associated with lysosomal enlargement and cell toxicity implicated in depression^8, 47^ and coronary artery disease^48^. A further locus on chromosome 5 is in an intergenic region. Finally, a locus on chromosome 3 located in *Myosin Heavy Chain 15*, a gene coding for a class of motor proteins that is highly expressed in the brain in humans. Our GWAS is considerably better powered than previous anxiety GWAS, including 23 453 cases compared to 3 695 cases^11^, in addition to having a homogenous study design for all UK Biobank analyses. While we did not see evidence for replication in two smaller anxiety cohorts, two of the five loci we identify had significant evidence for formal replication in larger independent samples for the related phenotypes of trait neuroticism and major depressive disorder. Phenotypic and genetic correlations of these phenotypes with anxiety are substantial.

Gene level analyses of Lifetime Anxiety Disorder confirmed the association with *Neurotrophic Receptor Tyrosine Kinase 2 (NTRK2*; p = 1.79×10^−6^) and *Transmembrane Protein 106B (TMEM106B;* p = 6.28×10^−10^) and enabled the identification of several additional protein coding genes associated with anxiety disorder. Specifically, *Phospholipase C Gamma 1* (*PLCG1*; p = 2.96×10^−7^), *Ring Finger and WD Repeat Domain 2 (RFWD2*, p=4.60×10^−7^*), Zinc Fingers and Homeoboxes 3 (ZHX3* p=5.21×10^−7^), *Glutamate Decarboxylase 2* (*GAD2*; p = 7.00×10^−7^), and *Lipin 3* (*LPIN3;* p = 2.54×10^−6^). *NTRK2* has been implicated in several neuropsychiatric traits and disorders including emotional arousal^38^, autism^39^, suicide^40–42^, Alzheimer’s disease^43^, alcohol dependence^46^, and treatment response^44–45^, although many of these are candidate gene studies and are thus less likely to represent robust findings. *TMEM106B* was recently found to be associated with both a broad depression phenotype and probable major depression in the UK Biobank^47^, and with major depression in the Psychiatric Genomic Consortium meta-analyses^8^. This is unsurprising given the phenotypic and genetic overlap between anxiety and depression^10^. *TMEM106B* is widely expressed throughout normal human cell types and tissue, including the fetal brain, and adult frontal cortex. A recent investigation demonstrated that a noncoding variant on chromosome 7 (rs1990620) interacts with *TMEM106B* via long-range chromatin looping interactions, mediating increased *TMEM106B* expression, which is strongly associated with lysosome enlargement and cell toxicity^49^. Conditional analyses indicate that the locus associated with Lifetime Anxiety Disorder in the UK Biobank is the same locus which results in increased expression of *TMEM106B*, suggesting a role for lysosome enlargement and cell toxicity in anxiety (see **Supplementary Information, Figures 10-11)**.

*Glutamate decarboxylase 2* (*GAD2*) encodes for an enzyme that synthesises gamma-aminobutyric acid (GABA) from L-glutamic acid. GABA is the principal inhibitory neurotransmitter in the mammalian central nervous system and has well-established associations anxiety. Specifically, variation in *Glutamate decarboxylase* genes have been associated with internalising disorders^50^. Furthermore, GABAergic deficits are observed in patients with, and animal models of, anxiety and depression^51^. Finally, benzodiazepines, a class of pharmacological agent with strong anxiolytic effects, act through binding to a specific GABA receptor, facilitating GABAergic inhibitory effects^52^.

Several findings relating to the polygenicity of anxiety are notable. First, SNP-heritability (h^2^_SNP_) estimates for Lifetime Anxiety Disorder (h^2^_SNP_=25.7%) and Current Anxiety Symptoms (h^2^_SNP_=30.8%) in UK Biobank were especially large. Typically h^2^_SNP_ estimates are less than half twin h^2^ (40-60%)^4, 5^. Heritability estimates from twin studies take into account genetic influence of both common and rare genetic variants, whereas h^2^_SNP_ only takes into account the additive effect of common variants. Our results suggest that a large proportion of heritable variance in anxiety is attributable to common genetic variants, with the effects spread over many hundreds or thousands of loci ^53^. Our h^2^_SNP_ estimates are also much higher than those derived from previous studies of anxiety^11, 22^, which may reflect the homogeneity of the UK Biobank sample in terms of assessment, sampling, and genotyping.

We also explored genetic overlap between anxiety phenotypes. These suggest that common genetic effects on anxiety are shared across different levels of severity, as well as across varying samples and definitions, and with both neuroticism and depression. Polygenic scores derived from primary phenotype predicted ∼0.5% of the variance in Lifetime Anxiety Disorder case control status. We note that the moderate to high h^2^_SNP_ for the UK Biobank anxiety phenotypes suggests a PRS for anxiety will improve as larger discovery GWAS samples are used for the creation of PRS, increasing the signal to noise ratio in the construction of polygenic scores.

Genetic overlap was detected between anxiety and a broad range of other traits. Significant positive genetic correlations were detected between Lifetime Anxiety Disorder and several other psychiatric phenotypes (schizophrenia, bipolar disorder, insomnia, ADHD and cross-disorder analyses), indicating a degree of shared genetic risk between psychiatric traits more generally. Interestingly, given some epidemiological evidence for an association between these phenotypes^54^, a significant genetic correlation was observed between anxiety and coronary artery disease. This may reflect the previously known epidemiological^55^, and genomic overlap between major depressive disorder and coronary artery disease^8^. We interpret these findings in the context of several limitations. There is substantial sample overlap within the two UK Biobank anxiety phenotypes and between UK Biobank anxiety and several of the other phenotypes which also use the UK Biobank sample. However, LD score regression is robust to sample overlap^56^, so genetic correlations are unlikely to be biased by this.

Although the intergenic genome-wide significant loci on chromosome 9 previously associated with neuroticism^36, 37^ and depression^8^ replicates in the combined Lifetime Anxiety Disorder meta-analysis, the significance is attenuated. Nonetheless, this provides good evidence for a role of this region in lifetime anxiety. Not all genome-wide significant SNPs observed in the individual cohorts survive the combined sample Lifetime Anxiety Disorder meta-analysis. This lack of replication may be due to a winner’s curse effect^57^, the relatively modest power gain from the additional samples, or phenotypic and genetic heterogeneity across the studies. As noted above, the UK Biobank is a highly homogenous group of individuals, all based within the UK with the same sampling and phenotyping instrument used. The iPSYCH sample is also highly homogenous group of individuals, all based within Denmark. Conversely, the ANGST study is comprised of many smaller samples, using different GWAS arrays and varied phenotyping from across multiple continents. Thus, it is possible that sample heterogeneity contributes to the non-replication of some of our GWAS significant loci. Given that our analyses suggest anxiety is a highly polygenic trait, driven by small effects across many SNPs, it is likely that it remains somewhat underpowered, despite the sample being substantially larger than any prior genome-wide study of anxiety to date.

Although homogeneity within UK Biobank may be a significant factor in the success of our analyses and the high heritabilities found, this sample homogeneity is itself a limitation, as our analyses were thus limited across all three samples to individuals of European ancestry^58^. Another limitation is that the Lifetime Anxiety Disorder phenotype relies both on retrospective recall by the subject and accurate diagnosis by an unknown clinician, each of which are likely to contain error. However, there are high genetic correlations between this category and every other anxiety phenotype assessed, suggesting the category of Lifetime Anxiety Disorder has utility.

This study used the largest currently available dataset to investigate the role of common genetic variation in anxiety. Although no sufficiently powered studies are available to enable strict statistical replication^59^, the study provides the first well powered characterisation of the common genetic architecture of anxiety. Analyses demonstrate that a large proportion of the broad heritability of severe and pathological anxiety is attributable to common genetic variants, and that this is shared between anxiety phenotypes and other closely related psychiatric disorders. Several genomic regions, which include loci with known associations with internalising disorders were found to be significantly associated with anxiety in these analyses. These findings are likely to be of interest for targeted characterisation of the underlying biology of anxiety.

## Acknowledgements

This research has been conducted using the UK Biobank Resource, under application 16577.

This study represents independent research part funded by the National Institute for Health Research (NIHR) Biomedical Research Centre at South London and Maudsley NHS Foundation Trust and King’s College London. The views expressed are those of the author(s) and not necessarily those of the NHS, the NIHR or the Department of Health. High performance computing facilities were funded with capital equipment grants from the GSTT Charity (TR130505) and Maudsley Charity (980).

CK currently receives salary support from National Institute for Health Research (NIHR) and has previously received salary support from the Novo Nordisk UK Research Foundation, NIHR Biomedical Research Centre for Mental Health at South London and from the Maudsley National Health Service (NHS) Foundation Trust in the past. CR is supported by a grant from Fondation Peters to TE and GB. JH is supported by the National Institutes of Health grant R01 MH113665.

The iPSYCH team acknowledges funding from the Lundbeck Foundation (grant no R102-A9118 and R155-2014-1724), the Novo Nordisk Foundation for supporting the Danish National Biobank resource, and grants from Aarhus and Copenhagen Universities and University Hospitals, including support to the iSEQ Center, the GenomeDK HPC facility, and the CIRRAU Center.

T.C. Eley is part funded by a program grant from the UK Medical Research Council (MR/M021475/1).

K.L.P acknowledges funding from the Alexander von Humboldt Foundation and the UK Medical Research Council (MR/M021475/1).

## Author contributions

KP, TE, MH, KKN and GB conceived the study. KP, JC, SMM, CR, HG and SWC performed statistical analyses. CH, CK, HG, JC and KP performed phenotype and data QC for the UKBB samples. MM supervised the pre and post GWAS analysis pipeline for the iPSYCH sample. OM, MN, MBH, JBG, PBM, TW, DMH and ADB provided and processed samples for the iPSYCH sample. KP, JC, TE, GB wrote the manuscript. MH, KD, JH, JD, AM, MM gave advice and feedback at several stages of data generation and manuscript writing. All authors reviewed the manuscript.

## Supplementary Information

### Supplementary methods

#### Phenotype Creation in the UK Biobank

##### (1) Any anxiety diagnosis

Participants self-reported receiving a professional diagnosis of one or more of the following anxiety disorders: Social anxiety; any other specific phobia (e.g. disabling fear of heights, or spiders); panic attacks; anxiety, nerves or generalised anxiety disorder or agoraphobia. Participants were excluded from this analysis if they reported a professional diagnosis of any of the following: schizophrenia; bipolar disorder; attention deficit hyperactivity disorder; autistic spectrum disorder; eating disorder.

##### (2) Probable lifetime generalised anxiety disorder (GAD)

Participants were assessed as having probable generalised anxiety disorder if they met the following criteria:

i. Report having worried more than most people would in the same situation **OR** have had stronger worry than most people during their worst period of anxiety. **AND**
ii. Have had at least one time in their lives where they have felt worried, tense, or anxious most of the time for at least a month **AND**
iii. This frequent and persistent worrying has persisted 6 months or more **AND**
iv. Report having worried most days during their worst period of anxiety **AND**
v. **EITHER** had many different worries on their mind at the same time during the worst of their **OR** worried about more than one thing during the worst period of anxiety. **AND**
vi. Found it difficult to stop worrying **OR** were often unable to stop worrying **OR** often found themselves unable to control their worry. **AND**
vii. Suffered 3 or more somatic symptoms of anxiety during the worst period of anxiety (these include: feelings of restlessness; feeling easily tired; having trouble falling or staying asleep; feeling keyed up or on edge; increased irritability; experiencing tense, sore or aching muscles; having difficulty concentrating) **AND**
viii. Their anxiety resulted in significant impairment in roles or normal function **AND**
ix. They were not already classified as anxiety cases for the “Lifetime Anxiety Disorder” analysis

##### (3) Current anxiety symptoms

The 7 item Generalised Anxiety Disorder 7 (GAD-7) ^1^ screening tool was administered to 157,366 participants who took part in the UK Biobank online mental health follow up. This measure assesses generalised anxiety symptoms (such as inappropriate worry) that have been experienced over the preceding two-week period.

Where participants missed more than 2 items on the GAD-7, they were excluded from analyses. If participants missed one or two items, the score for the missing item/s was imputed as the average score for all completed items. A total score was then created for all participants and used in subsequent analyses as a quantitative phenotype.

Participants who scored 10 or more on the GAD-7 total score were considered to have “current anxiety symptoms”, and were used as cases for these subsequent analyses. This is a typical cut off point for some symptom severity using this screening measure ^1^. For sensitivity analyses exploring the dimensionality of anxiety symptoms, participants scoring between 5 and 10 on the GAD-7 total score were classed as having mild current GAD symptoms, scoring between 10 and 15 were classed as having moderate current GAD symptoms, and 15 or higher were classed as having severe current GAD symptoms. See Supplementary Table 1 for the total sample for each of these analyses.

##### (4) GAD-7 total score

126,443 individuals of western European descent completed the GAD-7 screening tool. Of these, 50% obtained a total score of 0, resulting in an extreme positive skew of this measure that was not easily corrected. This measure did not conform to assumptions of normality required for the association analyses undertaken in GWAS, and thus it was considered that any secondary analyses resulting from this would not be interpretable, and analyses treating GAD-7 as a continuous measure were not included.

##### (5) Screened Controls

All case/control phenotypes were compared against a subset of screened healthy controls. Healthy controls were screened for any self-reported or probable disorder based on their responses to the mental health online follow up. In addition, healthy controls did not report seeking any treatment for any psychiatric disorder, and were not prescribed psychotropic medication that might be indicative of a psychological disorder.

#### Brief description of the iPSYCH cohort and analyses

As described this includes anxiety cases and controls from the Lundbeck Foundation Initiative for Integrative Psychiatric Research (iPSYCH; http://ipsych.au.dk) that were specifically selected to match the same anxiety inclusion and exclusion criteria used in the present UK Biobank analysis. Genotypes for the primary Denmark cohort came from Guthrie cards held by the Danish Neonatal Screening Biobank at Statens Serum Institut. Samples from this biobank are linked to the Danish register system via the unique Danish personal identification number. The individuals were born between 1981 and 2005 and had to be alive and a resident of Denmark on their first birthday and have a known mother. Specifically, this analysis included those cases who met criteria for lifetime diagnosis of one of the five core anxiety phenotypes and did not meet criteria lifetime diagnosis of schizophrenia; bipolar disorder; autistic spectrum disorder; attention deficit hyperactivity disorder; or eating disorders (n = 2829), and controls (n = 10,386). Diagnoses were assigned by psychiatrists at inpatient and outpatient psychiatric services, and were identified using the Danish Psychiatric Central Research Register ^2^. Controls were randomly selected from the same nationwide birth cohort. The DNA from the dried blood spots was extracted, whole-genome amplified in triplicates and genotyped in 23 batches based on birth year. The first wave (batch) contains the youngest participants (born in 2004) and wave 23 consists of the oldest participants (born in 1981).

Genotyping was performed on Illumina’s PsychChip array at the Broad Institute of MIT and Harvard. GenCall (Akhunov et al. 2009) and Birdseed ^3^ were used to call variants with MAF > 0.01. Callsets were merged after pre-QC on individual call sets. Data processing and GWAS analyses were performed on secure servers at the GenomeDK high-performance computing cluster (http://genome.au.dk).

DNA preparation, genotyping, genotype calling, quality control methods, and imputation were performed on the broader iPSYCH cohort waves, which also included psychiatric cases with non-AN diagnoses (schizophrenia, depression, anorexia nervosa, bipolar disorder, autism spectrum disorder, and attention-deficit/hyperactivity disorder). The full iPSYCH cohort described contains ∼86,000 individuals, including about 57,000 cases with at least one of the noted psychiatric disorders and around 30,000 controls who do not have any of the noted psychiatric disorders that are investigated in iPSYCH. Each wave that was processed had ∼3,500 participants. More information about the iPSYCH study can be found here ^4^. Ancestry principal components and principal components that captured batch effects and other possible sources of variation were included as covariates.

#### Additional Genotype and Sample QC

Several QC steps were taken in addition to the UK Biobank QC pipeline.

##### Sample QC

Individuals were identified as biologically male if they had X-chromosome heterogeneity > 0.9, and as biologically female if they had X-chromosome heterogeneity > 0.5. Individuals were excluded from analyses if there was a discrepancy between their self-reported and biological sex based on these thresholds (401 individuals removed).

One member of any related pair was removed from further analyses (KING ≥ 0.044).

Individuals were also removed if overall genotype missingness was ≥ 0.02. Analyses were restricted to individuals of European descent as assessed through 4 means clustering (n = 462,065).

Genomic analyses were restricted to individuals of Western European descent as identified by 4 means clustering, based on the top principal components derived from the whole sample.

##### Genotype QC

In addition to UK Biobank genotype QC, variants were removed if HWE ≤ 1×10^−6^, GENO ≤ 0.02.

#### SNP heritability estimates

Conversion of the point estimate of heritability derived from BOLT-LMM ^5^ to the liability scale was undertaken as per procedures described in Lee et al. 2012 ^6^. Standard errors were converted to variances estimates, and converted to the liability scale as per procedure described in Lee et al. (2011) ^7^.

#### Gene-wise analysis

Gene based enrichment analyses were used to consider possible associations between “Lifetime Anxiety Disorder” at the aggregate gene level that may not be detectable at the SNP level in highly polygenic traits. MAGMA performs multiple regression on biologically informed gene sets with the outcome trait, while accounting for LD structure between SNPs^8^. Gene-sets were annotated according to proximity, including the gene region +/− 500 Mb.

#### Polygenic prediction of anxiety

Polygenic scores were created for each target sample using clumping to obtain SNPs in linkage equilibrium with an r^2^ < 0.1 within a 2500 kb window. Polygenic scores were created using all remaining SNPs at each of eight different p-value thresholds (0.001, 0.05, 0.1, 0.2, 0.3, 0.4, 0.5, 1). Logistic regression models were run in PRSice-2^9^ to assess the association between polygenic scores created at each threshold and “Lifetime Anxiety Disorder” case/control status, again with the same controls as in the main analysis. Ten thousand permutations were run for each subgroup to derive an empirical p-value robust to multiple testing.

#### Partition heritability

SNP heritability was partitioned using the “baseline model”, consisting of 53 functional categories, fully described elsewhere ^10^.

### Supplementary results

#### Partitioned heritability

There was a significant enrichment of SNP heritability for “Lifetime Anxiety Disorder” in regions of the genome known to be highly conserved across 29 Eutherian mammals ^30^, with 2.6% of SNPs accounting for 34% of the heritability (*se* = 0.08; *Enrichment metric* 13.01, *Enrichment se* = 3.01, *Enrichment p* = 0.0001, coefficient p = 6.7×10^−05^). See Supplementary Figure 8 for a bar plot showing relative enrichment of the SNP h^2^ across the 53 baseline genomic regions tested.

#### Exploration of TMEM106B gene locus

Steps were taken to ascertain whether the locus on chromosome 7 associated with “Lifetime Anxiety Disorder” is part of the same locus as variant rs1990620 previously found to play a putatively causal role in downstream expression of the TMEM106B and related cellular damage in frontotemporal degeneration ^11^. (1) A region plot of the TMEM106B locus was created tagging rs1990620 to establish whether this SNP is in high linkage disequilibrium with rs3807866. (2) Associations analyses were performed using SNPtest 32 conditioning on the putatively causal variant rs1990620 and (3) on the index variant rs3807866 to test whether these SNPs were independently associated with “Lifetime Anxiety Disorder” in the UK Biobank, or part of the same locus.

The putatively causal variant, rs1990620, identified by Gallagher et al. 2017 was found to be in high linkage disequilibrium with the index variant rs3807866 annotated to the TMEM106B gene associated with “Lifetime Anxiety Disorder” (see Supplementary Figure 9). When analyses of this region on chromosome 7 were adjusted for the effects of either this variant or the index variant identified in the “Lifetime Anxiety Disorder” genome wide association (rs3807866), no association with the “Lifetime Anxiety Disorder” phenotype remained (see Supplementary Figure 10). This indicates that the locus identified by Gallagher et al., 2017 is, in essence, the same locus which we find.

### Supplementary Figures

**Supplementary Figure 1.**
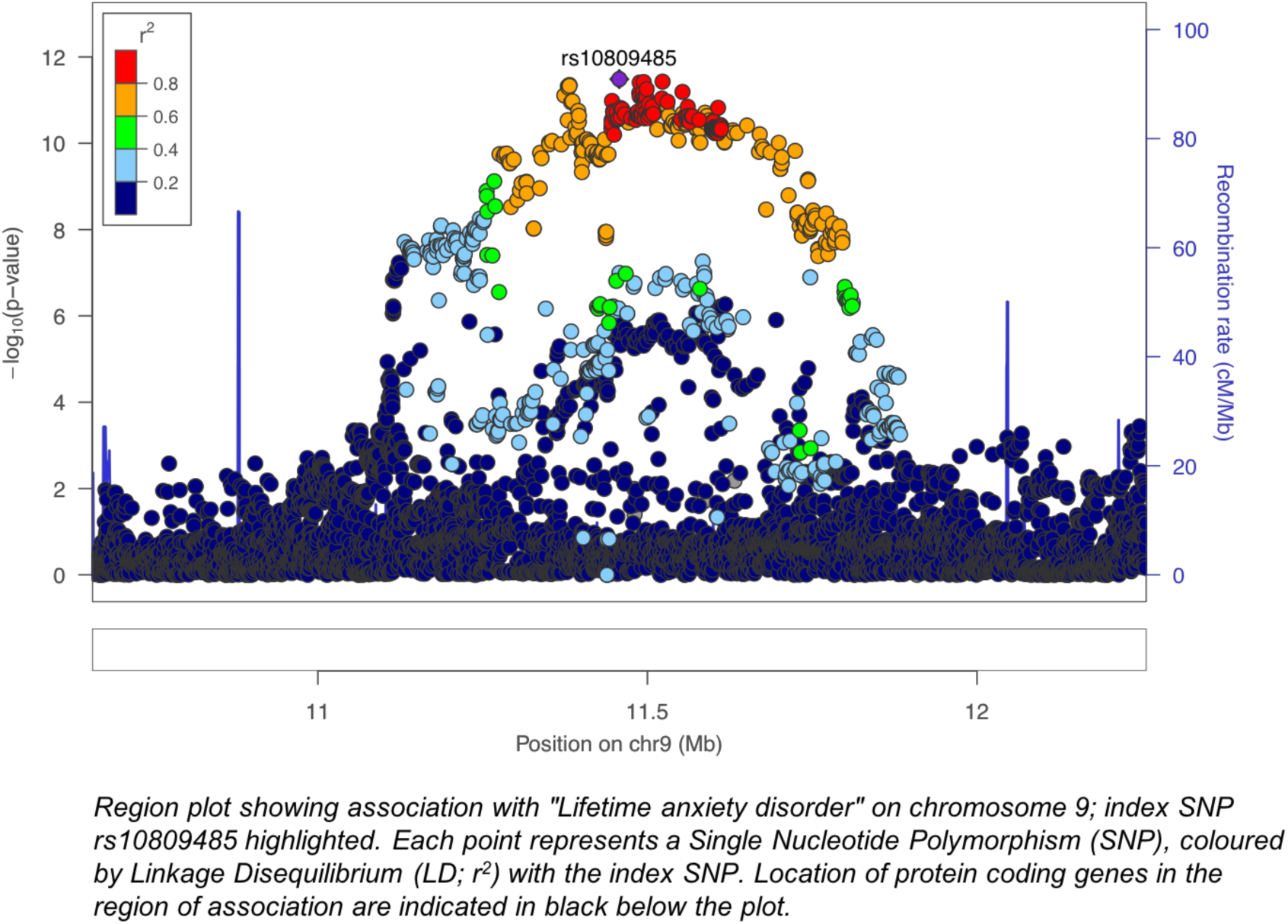
Region plot of rs10809485 locus in “lifetime anxiety disorder” genome-wide analyses

**Supplementary Figure 2.**
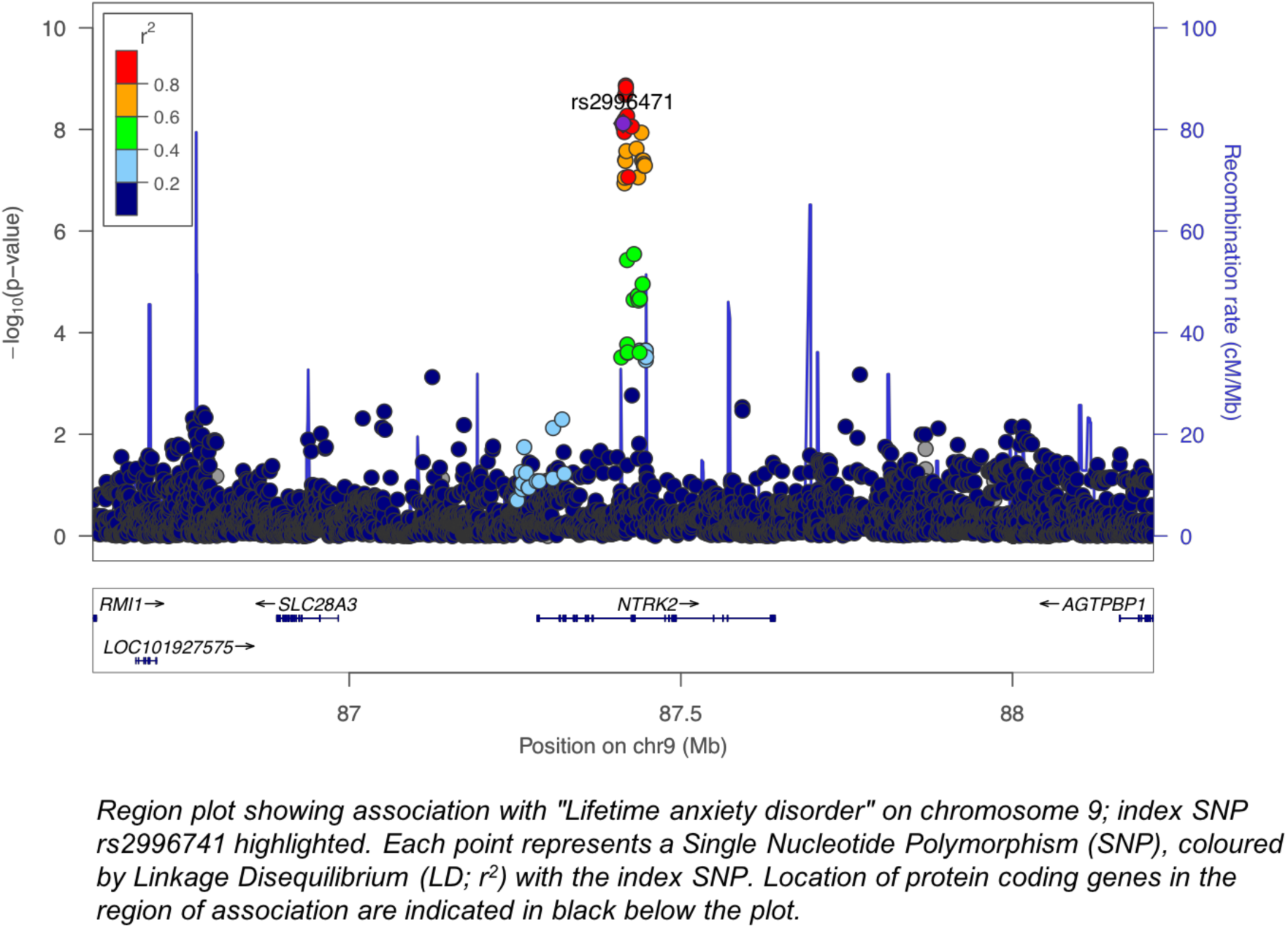
Region plot of rs2996471 locus in “lifetime anxiety disorder” genome-wide analysis

**Supplementary Figure 3.**
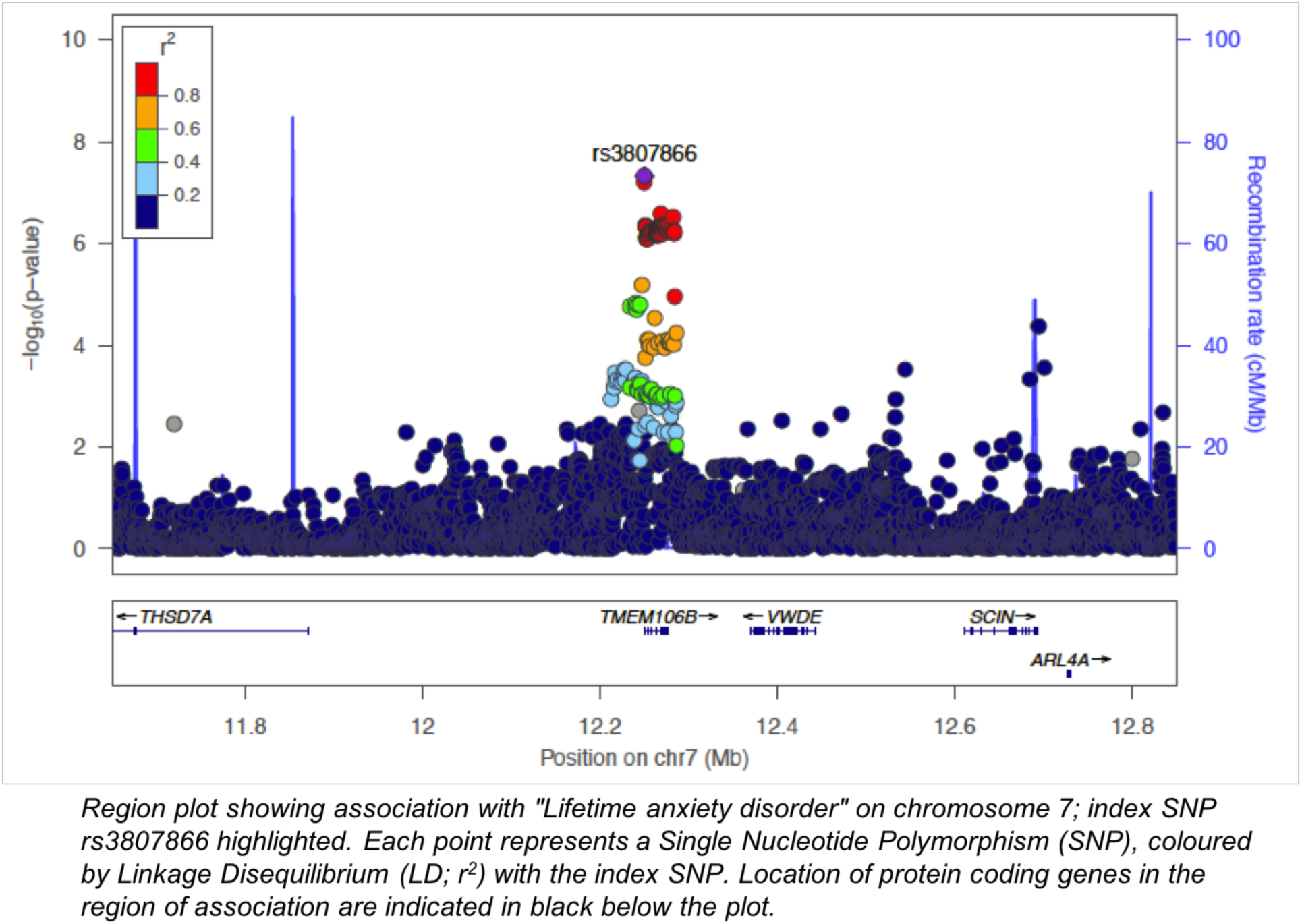
Region plot of rs3807866 locus in “lifetime anxiety disorder” genome-wide analysis

**Supplementary Figure 4.**
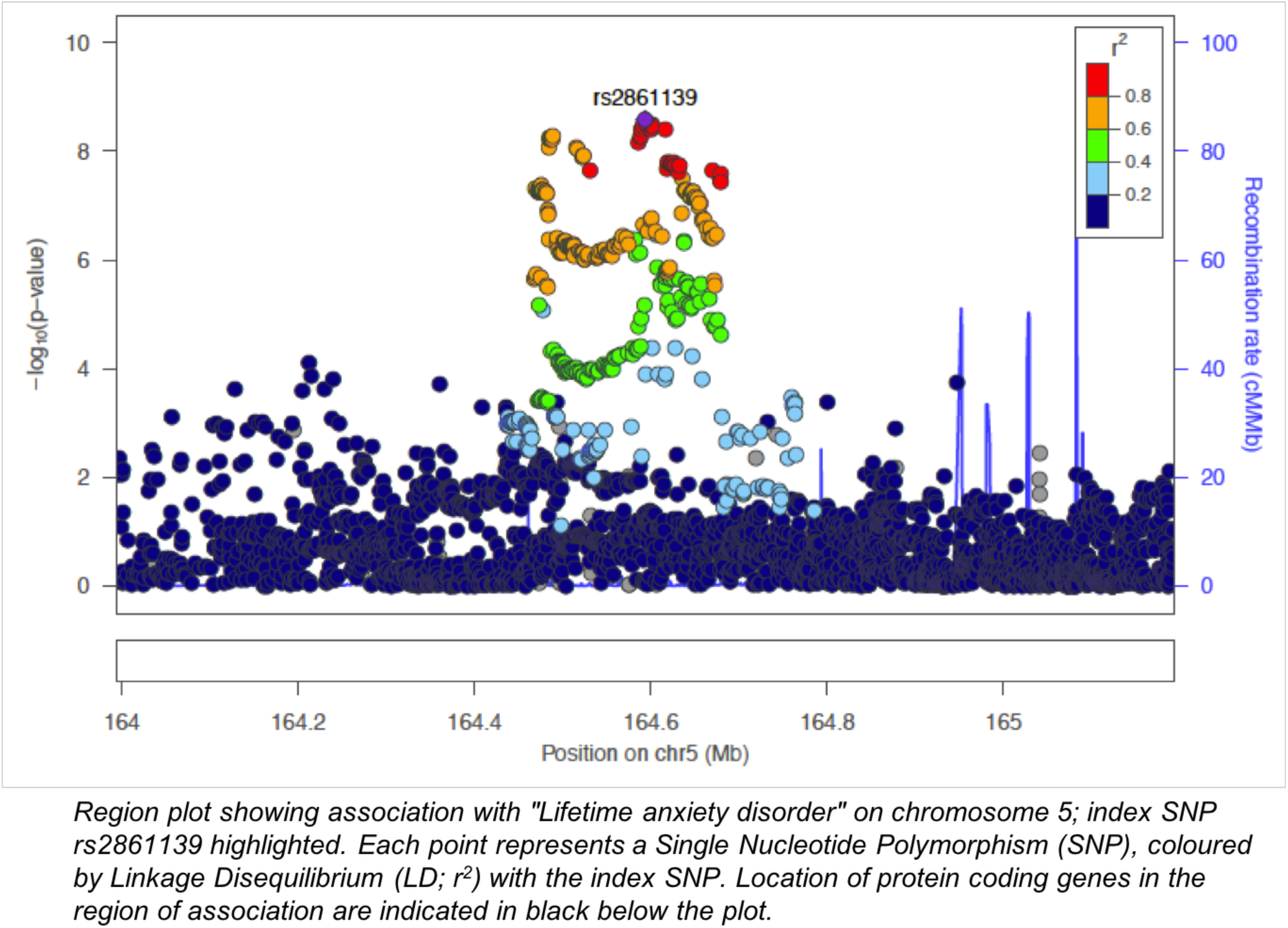
Region plot of rs2861139 locus in “lifetime anxiety disorder” genome-wide analysis

**Supplementary Figure 5.**
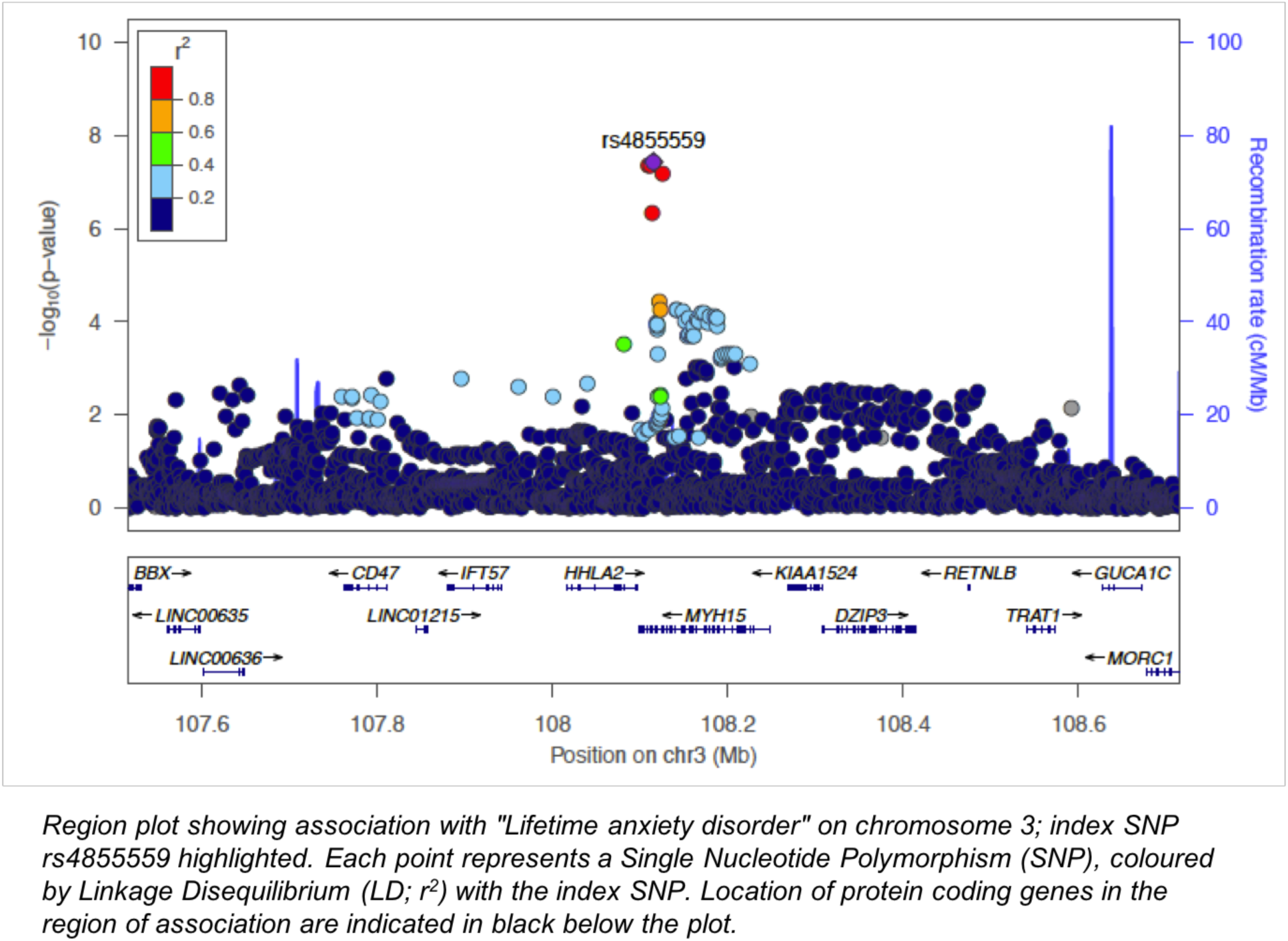
Region plot of rs4855559 locus in “lifetime anxiety disorder” genome-wide analysis

**Supplementary Figure 6.**
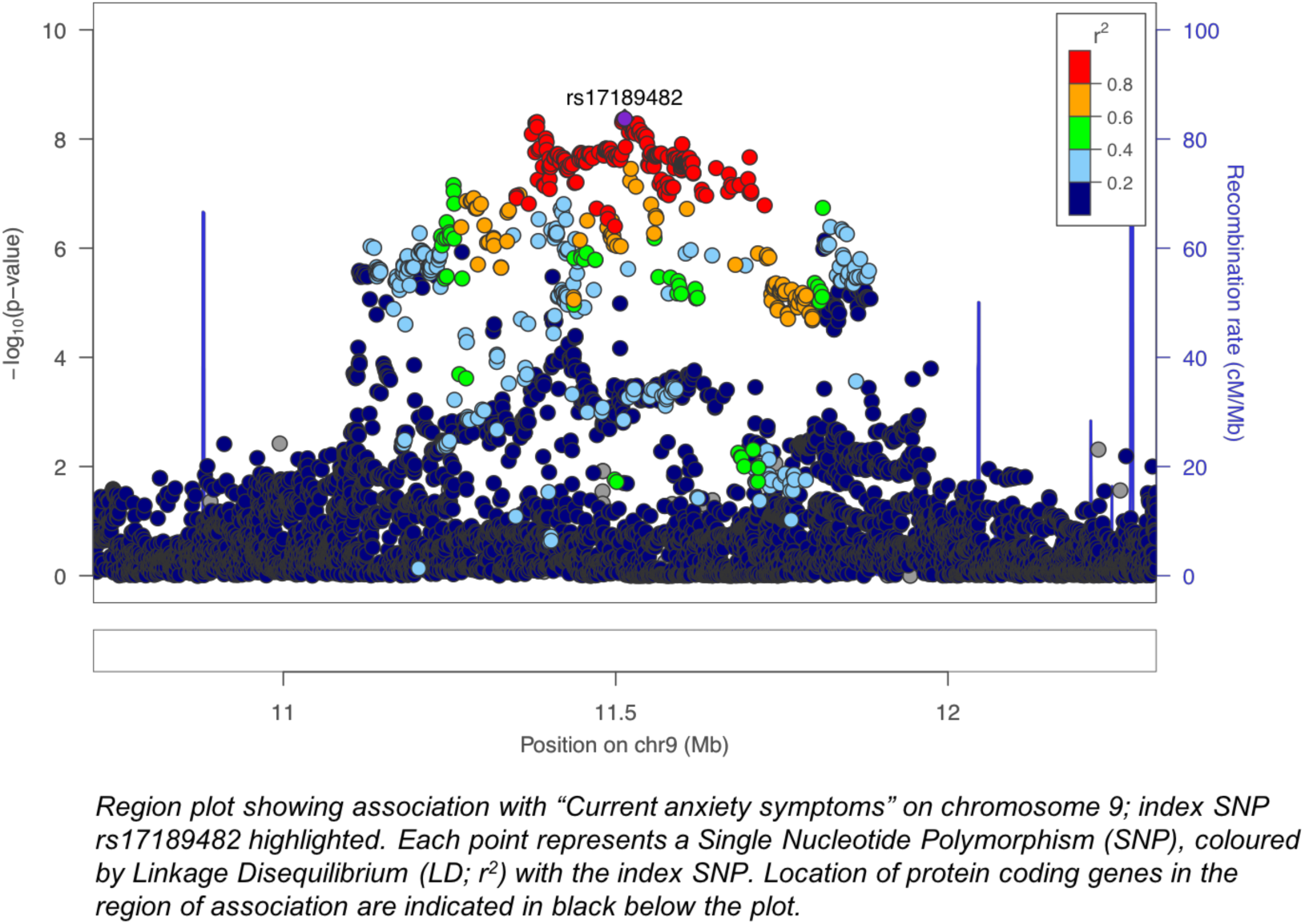
Region plot of rs10809485 locus in “Current anxiety symptoms” genome-wide analyses

**Supplementary Figure 7.**
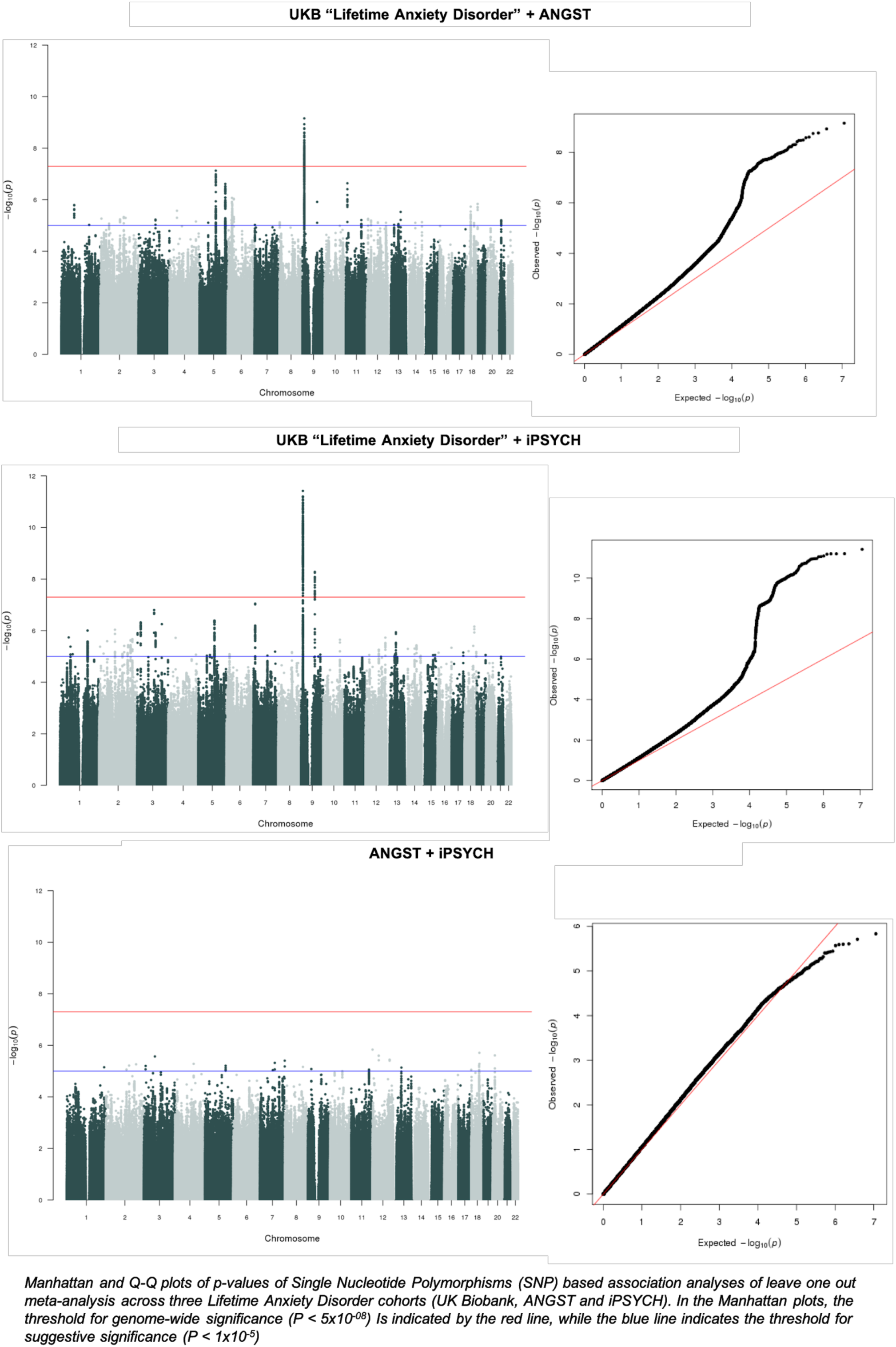
Leave One Out Manhattan and Q-Q plots for meta-analysis between all Lifetime Anxiety Disorder Samples

**Supplementary Figure 8.**
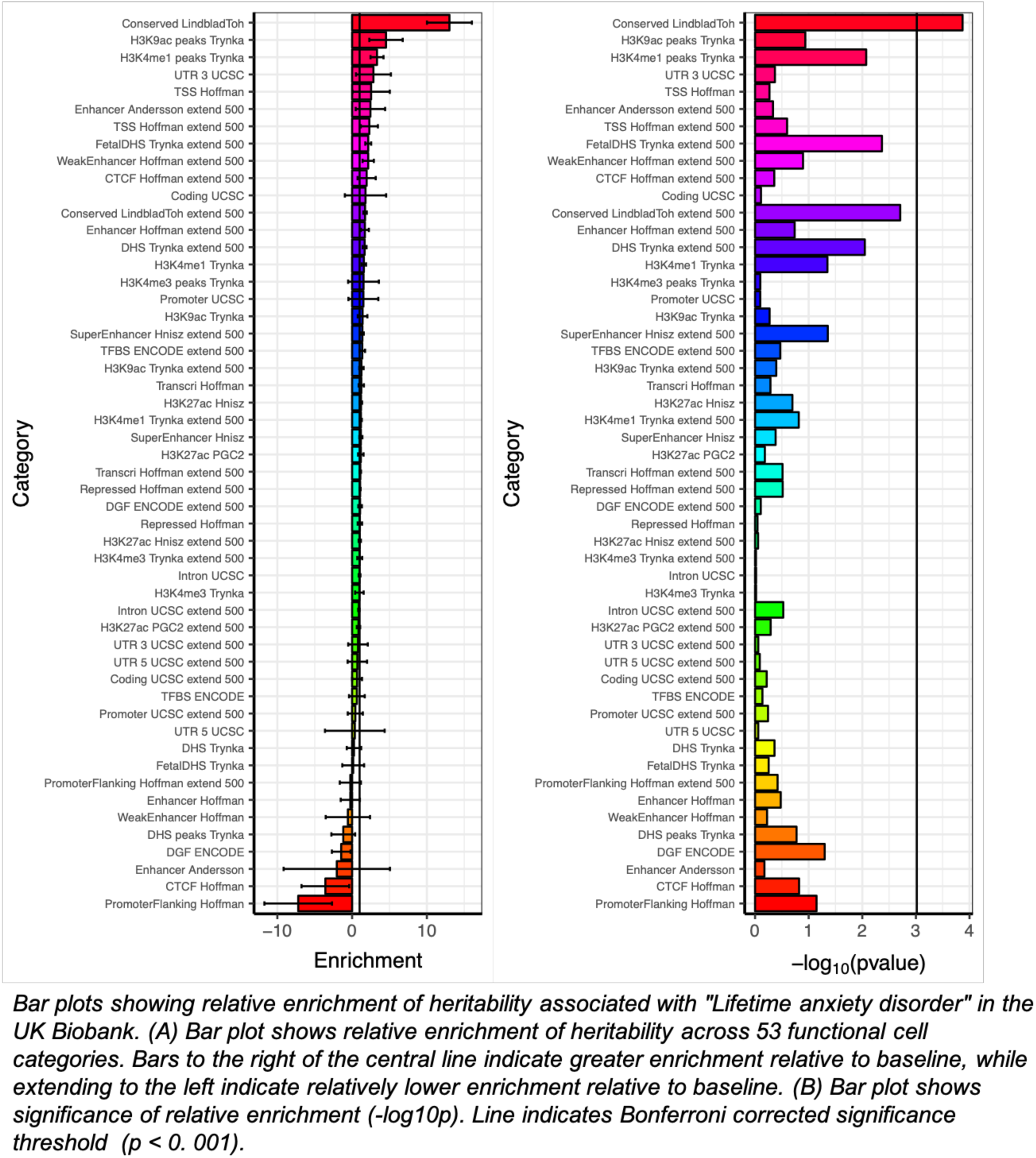
Bar plot showing relative enrichment of SNP heritability across functional categories for “Lifetime anxiety disorder” phenotype

**Supplementary figure 9.**
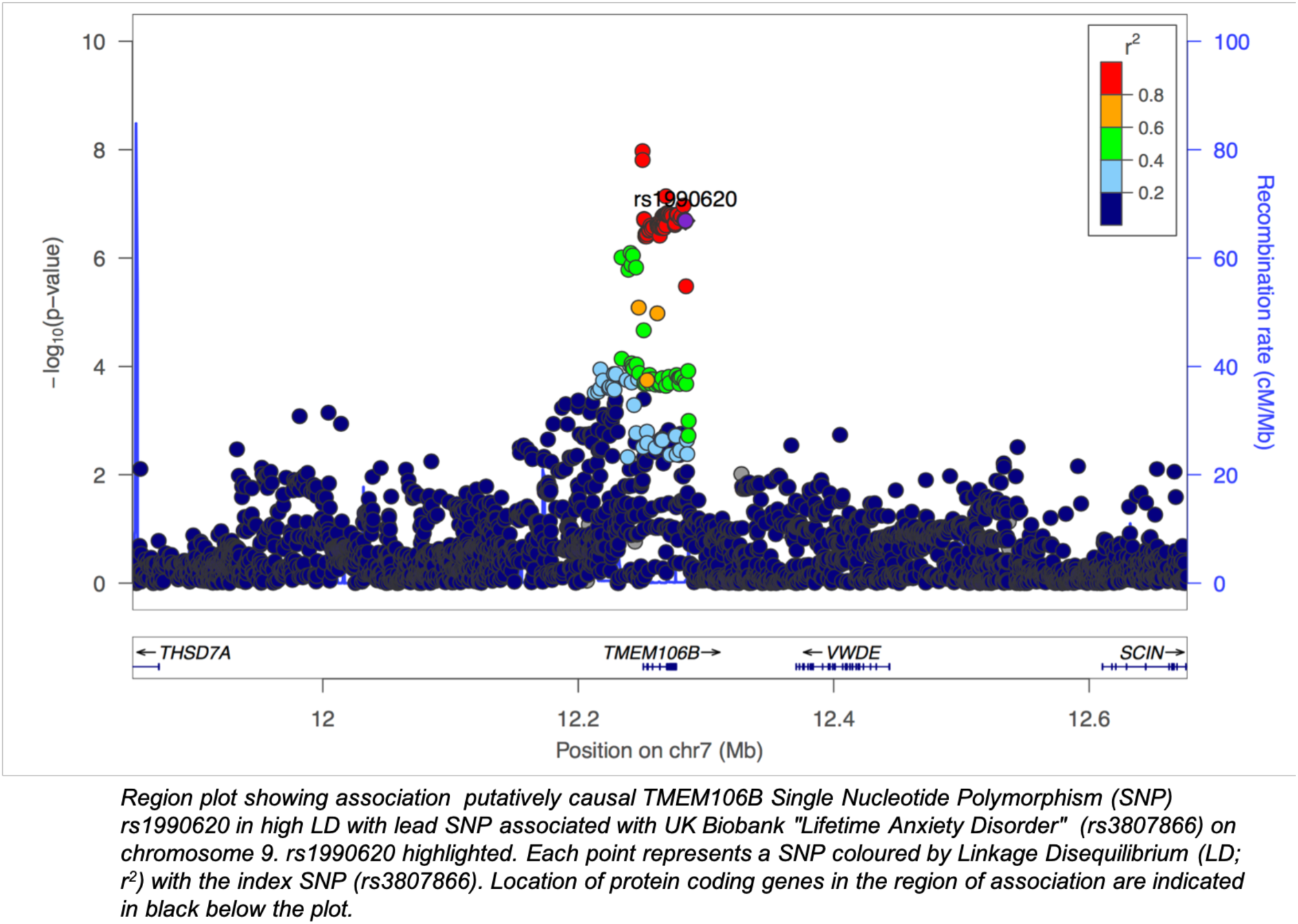
Region plot showing rs1990620 in high LD with lead variant associated with “Lifetime Anxiety Disorder” in the UK Biobank

**Supplementary Figure 10.**
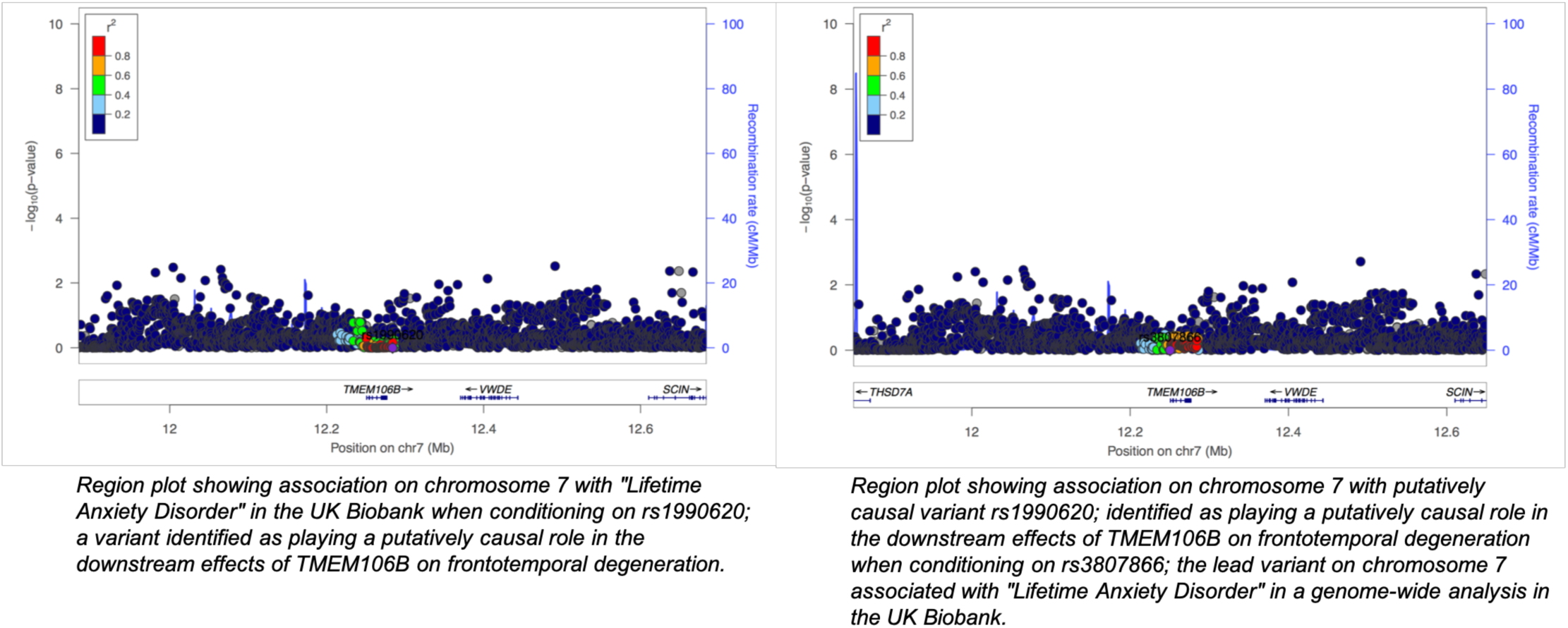
Region plots showing the effects of association in the TMEM106B gene locus when conditioning on the lead variant in UK Biobank “Lifetime Anxiety Disorder”, or a variant putatively part of the causal pathway for frontotemporal degeneration (rs 1990620)

